# The Zymoseptoria tritici avirulence factor AvrStb6 accumulates in hyphae close to stomata and triggers a wheat defense response hindering fungal penetration

**DOI:** 10.1101/2024.01.11.575168

**Authors:** Julien Alassimone, Coraline Praz, Cécile Lorrain, Agustina De Francesco, Cristian Carrasco-López, Luigi Faino, Lukas Meile, Andrea Sánchez-Vallet

**Affiliations:** Plant Pathology, Institute of Integrative Biology, ETH Zürich, Zürich, Switzerland; Centro de Biotecnología y Genómica de Plantas (CBGP/Universidad Politécnica de Madrid-Instituto Nacional de Investigación Agraria y Alimentaria/Centro Superior de Investigaciones Científicas (INIA/CSIC). Campus de Montegancedo. Pozuelo de Alarcón (Madrid) Spain.

**Keywords:** Plant resistance, wheat fungal pathogen, stomatal-mediated resistance, transcriptomics, gene-for-gene interaction, septoria tritici blotch

## Abstract

*Zymoseptoria tritici*, the causal agent of septoria tritici blotch, is one of Europe’s most damaging wheat pathogens, causing significant economic losses. Genetic resistance is a common strategy to control the disease, *Stb6* being a resistance gene used for over 100 years in Europe. This study investigates the molecular mechanisms underlying Stb6-mediated resistance. Utilizing confocal microscopy imaging, we identified that *Z. tritici* epiphytic hyphae mainly accumulates the corresponding avirulence factor AvrStb6 in close proximity to stomata. Consequently, the progression of AvrStb6-expressing avirulent strains is hampered during penetration. The fungal growth inhibition co-occurs with a transcriptional reprogramming in wheat characterized by an induction of immune responses, genes involved in stomata regulation, and cell wall-related genes. Overall, we shed light on the gene-for-gene resistance mechanisms in the wheat - *Z. tritici* pathosystem at the cytological and transcriptomic level, and our results highlight that stomata penetration is a critical process for pathogenicity and resistance.

## INTRODUCTION

Disease outcomes in plant-pathogen interactions are determined by the pathogen’s ability to infect a host and the host’s capacity to hamper the invader’s progression. This intricate interaction forces both organisms to gain novel mechanisms to coexist with each other through evolution (Sacristán and García-Arenal 2008; Krattinger and Keller 2016). The plant immune system protects against invading microorganisms through the activity of resistance genes that recognize specific components of pathogens, while pathogens adapt with numerous infection strategies to circumvent the immune response and successfully colonize the host (Sánchez-Vallet et al. 2018). Host detection of common pathogen molecular patterns triggers an immune response that frequently culminates in the arrest of pathogen progression (Jones and Dangl 2006; Cook et al. 2015; Kanyuka and Rudd 2019). This immune response is counteracted by invading microorganisms through the secretion of so-called effectors (Toruño et al. 2016). Plants have, in turn, evolved to specifically recognize particular forms of these effectors and hinder the pathogen progression. In these cases, effectors are known as avirulence factors, and they are specifically recognized by host resistance proteins in a gene-for-gene manner (Jones and Dangl 2006; Toruño et al. 2016). Although this type of resistance is broadly distributed in plant species, the mechanisms involved in arresting the growth of avirulent strains are frequently unknown.

The fast-evolving pathogen *Zymoseptoria tritici* is the causal agent of septoria tritici blotch (STB) and Europe’s most devastating wheat pathogen (Fones and Gurr 2015). STB disease is challenging to control due to the development of fungicide resistance by *Z. tritici* and the erosion of genetic resistance in modern cultivars (Brown et al. 2015; Torriani et al. 2015; O’Driscoll et al. 2014). Infection of *Z. tritici* is initiated by the germination of the asexual or sexual spores on the leaf surface. Then, the emerging hyphae penetrate the apoplast through stomata and wounds (Kema et al. 1996; Duncan and Howard 2000; Fones et al. 2017; Battache et al. 2022; Bernasconi et al. 2023). Intercellular hyphae of *Z. tritici* invade the leaf tissue without producing symptoms during the first phase of infection that lasts for a minimum of 7 days (Kema et al. 1996; Duncan and Howard 2000; Steinberg 2015). This asymptomatic phase is followed by a necrotrophic phase, which is characterized by the development of chlorotic and necrotic lesions and the production of pycnidia, the asexual reproductive structures (Sánchez-Vallet et al. 2015; Steinberg 2015).

Until now, 21 resistance genes against STB have been mapped to the wheat genome (Brown et al. 2015), three of them (*Stb6*, *Stb16q* and *Stb15*) have been cloned (Saintenac et al. 2018, 2021; Hafeez et al. 2023). These resistance genes generally confer resistance against *Z. tritici* strains that harbor the corresponding avirulence factors. The resistance gene *Stb6*, present in approximately 15% of wheat cultivars, is one of the most frequently used resistance genes in breeding programs (Chartrain et al. 2005; Arraiano and Brown 2006; Brown et al. 2015). Stb6 is a wall-associated kinase (WAK) that recognizes the fungal avirulence factor AvrStb6 (Zhong et al. 2017; Kema et al. 2018; Saintenac et al. 2018). In contrast to what has been described for other pathosystems, *Z. tritici* avirulence factor recognition does not lead to a hypersensitive response (Cohen and Eyal 1993; Kema et al. 1996; Battache et al. 2022). Penetration to the apoplast is crucial for wheat resistance against *Z. tritici*. The wheat resistance gene *Stb16q*, encoding for a cysteine-rich protein kinase, and recognition of the avirulence factor Avr3D1 prevent *Z. tritici* penetration through the stomata (Saintenac et al. 2021; Battache et al. 2022; Meile et al. 2023). In addition, it was recently demonstrated that stomatal closure is associated with Stb16q- and Stb6-mediated resistance (Battache et al. 2022; Ghiasi Noei et al. 2022) and that wounding partially enables infection by avirulent strains (Battache et al. 2022; Bernasconi et al. 2023). A better understanding of Stb6-mediated resistance mechanisms against *Z. tritici* invasion would provide us with novel tools to improve control methods against *Z. tritici*. In this study, we demonstrated that AvrStb6 is accumulated in hyphae penetrating the stomata where the progression of avirulent strains is hindered. We characterized the specific transcriptional reprogramming occurring in the host upon recognition of AvrStb6 making use of isogenic lines of *Z. tritici* expressing or not *AvrStb6* and the purified protein AvrStb6. We showed that AvrStb6 recognition triggers the induction of genes involved in stomatal closure regulation and a defense response associated with the pathogen’s incapacity to colonize and infect the plant.

## RESULTS

### Stomatal penetration is hindered in incompatible interactions

To investigate at which infection stage AvrStb6 recognition hinders *Z. tritici* progression, we monitored the colonization of Chinese Spring (with Stb6) leaves by fluorescently labeled strains that either harbor a virulent or an avirulent allele of *AvrStb6* (strains 3D7 and 1E4, respectively; Figure 1). Successful penetration events of the virulent strain were observed as early as 6 dpi (Figure 1A and G). At that stage, hyphae of both strains colonized the leaf surface and were frequently in contact with stomata. Hyphae from the virulent strain penetrated the substomatal cavity and colonized the stomata proximate intercellular space. At 10 and 14 dpi, hyphae of the virulent strain thoroughly colonized the apoplast and the substomatal cavity (Figure 1B and C). In contrast, the avirulent strain was incapable of entering the stomata and remained on the leaf surface at the three observed time points (Figure 1E and F). Only in extremely rare events, we observed severely limited hyphal growth of avirulent strains inside the leaves (Supplementary Figure S2). Overall, our results suggest that stomata act as a main barrier for AvrStb6-expressing avirulent strains and that AvrStb6 recognition occurs at or in close proximity to the stomata.

**Figure 1.**
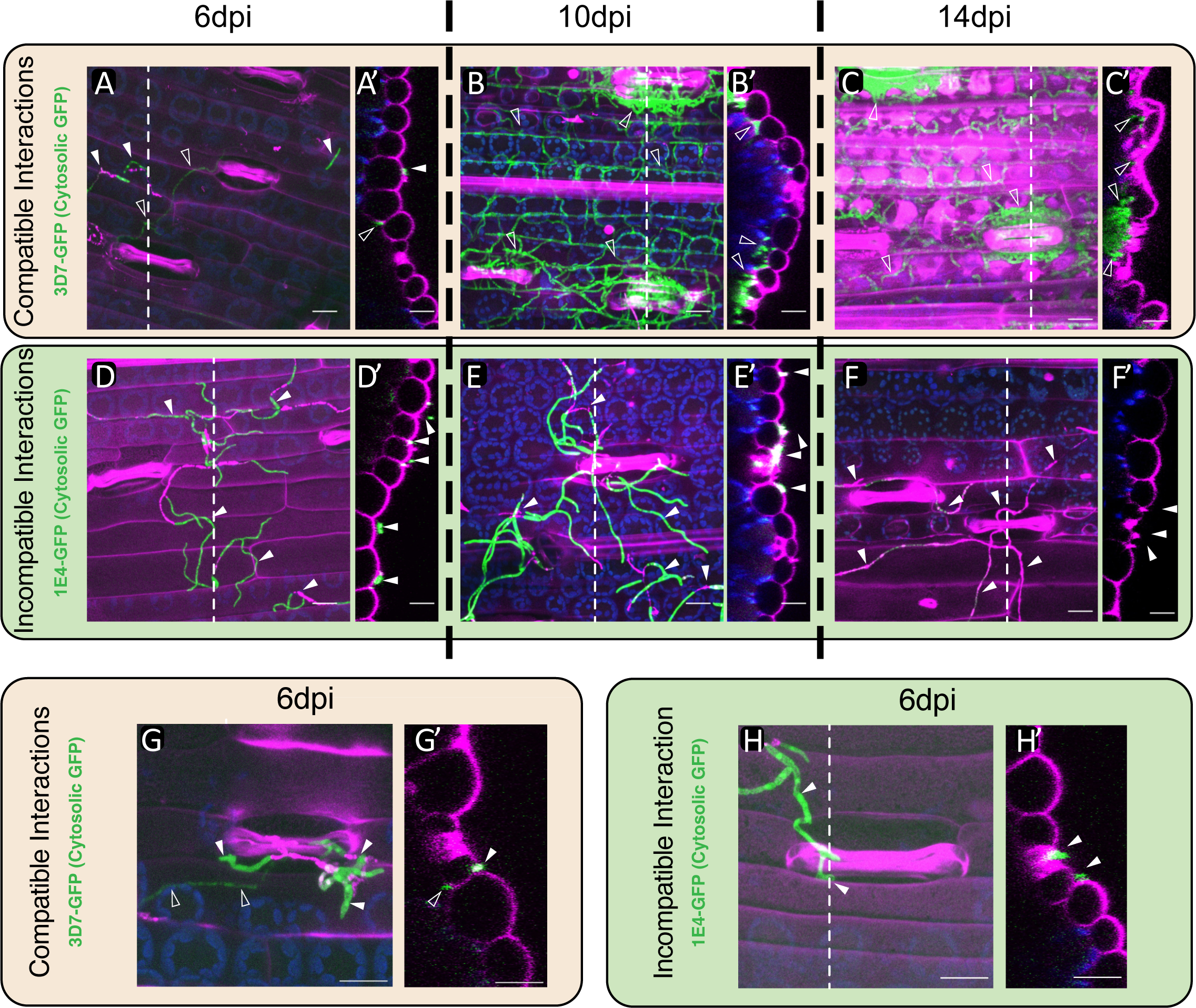
Recognition of AvrStb6 hinders the penetration of *Zymoseptoria tritici.* Confocal microscopy images of wheat cultivar Chinese Spring sprayed with the *Z. tritici* cytosolic reporter lines 3D7 (3D7-GFP; A-C and G) and 1E4 (1E4-GFP; D-F and H). **A**. At 6 days post-infection (dpi), the virulent strain 3D7 penetrated the leaf apoplast. **B.** At 10 and **C.** 14 dpi, 3D7 invaded the leaf apoplastic space and accumulated in sub-stomatal cavities. **D-F**. The avirulent strain remained at the leaf surface at all observed timepoints (6, 10, and 14 dpi; D, E and F, respectively). **G,H.** Stomata close-up views of compatible (G) and incompatible (H) interactions observed at 6 dpi. Images A-H are maximum projections, and A’-H’ are orthogonal views. Dashed lines indicate the location of the orthogonal view on the corresponding image. Images are overlays of GFP signal (green), chloroplast autofluorescence (blue) and plant cells stained with propidium iodide (pink). Full and empty white triangles highlight epiphyllous and apoplastic hyphae, respectively. Scale bar: 20 µm.

We corroborated the penetration capacity of avirulent strains by infecting wheat cultivar Chinese Spring with 3D7-GFP and a mutant line ectopically expressing the avirulent effector gene *AvrStb6*_1E4_. The lines expressing *AvrStb6*_1E4_ were impaired in stomatal penetration, while the control lines colonized the apoplast. These results demonstrate that AvrStb6 recognition prevents stomatal penetration (Supplementary Figure S3A-F). Additionally, to demonstrate the role of Stb6, we assessed stomatal penetration of the avirulent strain 1E4 in wheat near-isogenic lines (NILs) with and without *Stb6* in the Bobwhite background (Saintenac et al. 2018). Only *Stb6*-harboring wheat lines prevented penetration of 1E4 (Supplementary Figure S3G-L). These results demonstrated that AvrStb6-Stb6 interactions result in the pathogen being mostly unable to enter the leaf via the stomata.

### The avirulence factor AvrStb6 is expressed during stomata penetration and apoplast colonization

Although *AvrStb6* has been described to exhibit a peak of expression at the onset of the necrotrophic phase (Rudd et al. 2015; Zhong et al. 2017), its protein accumulation pattern at the cellular level remains undetermined. We used a 3D7 mutant expressing AvrStb6_1E4_-GFP to monitor the cell-specific accumulation of AvrStb6 during colonization of the wheat NILs (with or without *Stb6;* Figure 2 and Supplementary Figure S4). In addition, this *Z. tritici* reporter strain constitutively expresses cytosolic mCherry, allowing the visualization of hyphae colonizing wheat plants in microscopy assays. Regardless of the presence of *Stb6* in the host, epiphytic hyphae accumulated AvrStb6_1E4_-GFP mainly in hyphal cells in direct contact with stomata, and no signal was detected on the remaining hyphae at the leaf surface (Figure 2A, Supplementary Figure S4). In the compatible interaction, hyphae growing in the apoplast also exhibited a substantial accumulation of AvrStb6_1E4_-GFP (Figure 2B, Supplementary Figures S4B and D). These results indicate a cell-specific accumulation of AvrStb6_1E4_-GFP at the penetration sites and in the apoplastic space. This specific protein accumulation pattern is in accordance with the previously described tight *AvrStb6* gene expression pattern (Meile et al. 2018). During epiphytic growth, *AvrStb6* promoter activation was mostly restricted to fungal cells in direct contact with stomata and some of the hyphae extremities (Supplementary Figure S4K) (Meile et al. 2020). However, *AvrStb6* promoter activation was observed in all hyphae colonizing the apoplast (Supplementary Figure S4L) (Meile et al. 2020). These and our previously published results (Meile et al. 2020) demonstrate that transcription of *AvrStb6* is tightly regulated at the cellular level, and that the protein accumulation occurs on the site of *Z. tritici* colonization. We therefore hypothesize that the role of AvrStb6 is specifically related to the host invasion.

**Figure 2.**
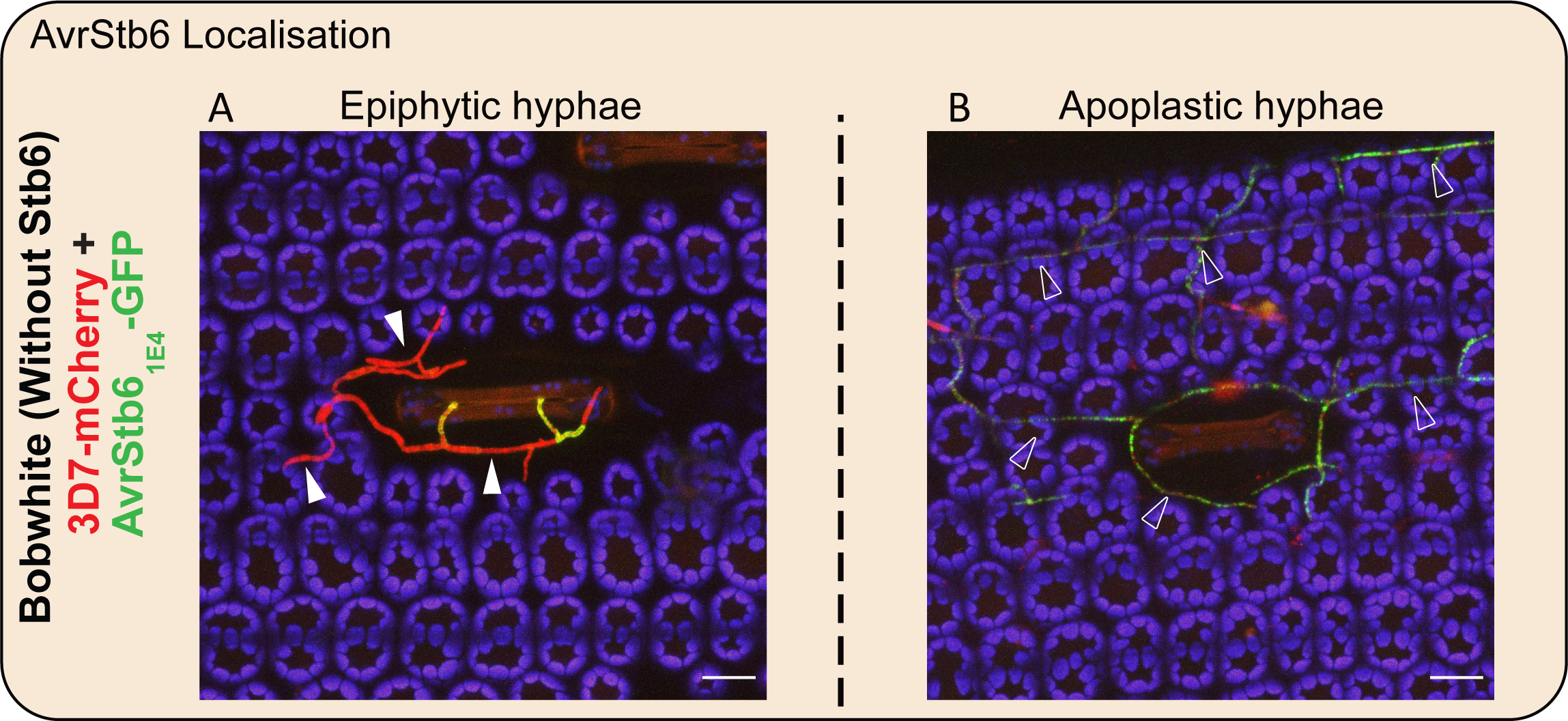
AvrStb6 is produced and accumulates at the infection site. **A-B.** Microscopy images of wild-type Bobwhite plants infected with the 3D7 mCherry-labelled *Zymoseptoria tritici* (red) strain expressing AvrStb6_1E4_ fused to GFP (AvrStb6_1E4_-GFP, green). Observations made at 6 days post-infection (dpi) of hyphae growing epiphytically (A) and in the apoplastic space (B). Images show GFP signal (green), mCherry signal (red) and chloroplast autofluorescence (blue). Images on A and B are maximum projections. Full and empty white triangles highlight epiphyllous and penetrated hyphae, respectively. Scale bar: 20 µm.

### *Z. tritici* AvrStb6 is a secreted protein

The avirulence factor AvrStb6 has a predicted signal peptide, suggesting that it is secreted (Brunner and McDonald 2018; Stephens et al. 2021) (Figure 3A). Interestingly, upon stomatal contact, AvrStb6_1E4_-GFP does not fully co-localize with the cytosolic reporter mCherry and seem to be located at *Z. tritici* cell surface or cell wall (Figure 3B). To test whether AvrStb6 is secreted from fungal cells and whether it remains bound to the fungal cell wall, we grew the 3D7 ectopic mutant lines expressing the GFP-tagged virulent and avirulent *AvrStb6* alleles (*AvrStb6*_3D7_-GFP and *AvrStb6*_1E4_-GFP, respectively) in axenic liquid media. We evaluated the presence of AvrStb6-GFP both in the spent medium and in the fungal cell by Western blot. As a control, we used a reporter 3D7 strain expressing cytosolic GFP. As expected, the cytosolic GFP was mainly present in the cell fraction (pellet). In contrast, both isoforms of AvrStb6-GFP (AvrStb6_1E4_-GFP and AvrStb6_3D7_-GFP) were predominantly present in the culture medium (Figure 3C). The protein detected in the pellet was minor, regardless of the treatment with urea, a cell disruptive agent. These results demonstrate that AvrStb6 has a functional signal peptide that enables the secretion of the effector protein and supports its extracellular function.

**Figure 3.**
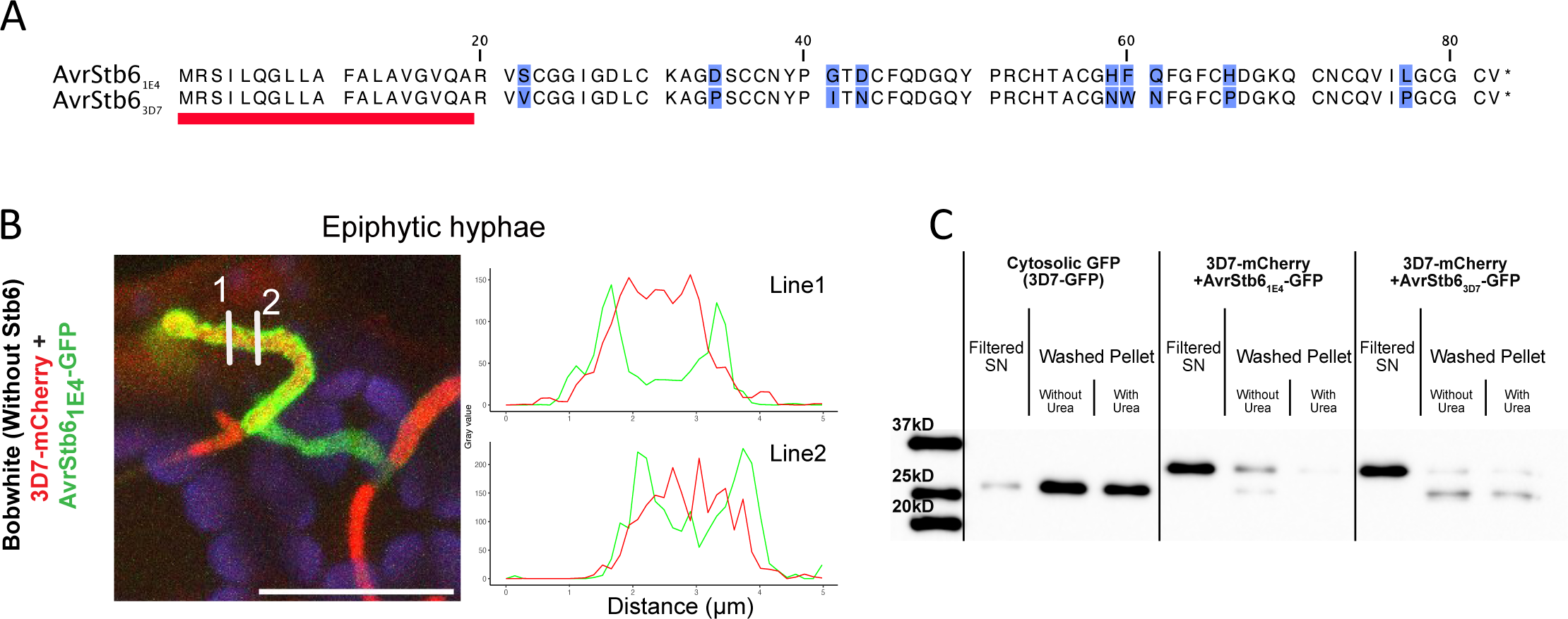
AvrStb6 is a secreted protein. . **A.** Protein alignment of 3D7 (virulent, AvrStb6_3D7_) and 1E4 (avirulent AvrStb6_3D7_) AvrStb6 protein isoforms of *Zymoseptoria tritici*. The red line indicates the predicted signal peptide. Blue boxes highlight amino acid polymorphism. **B.** Subcellular localization of AvrStb6_1E4_-GFP on epiphytic hyphae reaching a stomate of a wild-type Bobwhite plant infected with the 3D7 mCherry-labelled *Z. tritici* strain expressing AvrStb6_1E4_-GFP. Observations made at 6 days post-infection (dpi). Line charts indicate GFP (green) and mCherry (red) signal intensities (absolute grey value) distributed across the lines depicted in the corresponding image. Image is a maximum projection on the overlay of GFP signal (green), mCherry (red) signal, and chloroplast autofluorescence (blue). Observations made at 6dpi. Scale bar: 20 µm. **C.** Western blot analysis of *in-vitro* grown *Z. tritici* 3D7 strains expressing either a cytosolic GFP or each AvrStb6 isoform fused to GFP (3D7+AvrStb6_1E4_-GFP and 3D7+AvrStb6_3D7_-GFP). Fungal lines were grown in liquid media, and the western blot was performed on filtered supernatant (SN; secreted protein) and washed pellet (with and without urea treatment; non-secreted proteins, insoluble fraction) using an anti-GFP antibody. The expected molecular weights of GFP and the AvrStb6-GFP protein fusions are 26.94 and 36 kD, respectively.

### AvrStb6 triggers an immune response in resistant wheat cultivars

To determine the host transcriptomic response to AvrStb6 recognition, we compared the transcriptomic profiles of leaves infiltrated with a purified AvrStb6 protein (avirulent isoform) and of leaves infected with a strain of *Z. tritici* expressing the avirulent isoform of AvrStb6. We infiltrated the *Pichia-*produced avirulent isoform of AvrStb6 (Supplementary Figure S5) into leaves of cultivar Chinese Spring and performed RNA sequencing of the leaves after 1, 2, and 5 hours post-infiltration. Phosphate-buffered saline (PBS) infiltration was used as control. The purified avirulent isoform of AvrStb6 protein did not produce any visible symptoms in wheat leaves and was shown to be recognized by Chinese Spring since its infiltration prior to the infection with the virulent strain 3D7 led to a reduction of virulence (Figure 4A). In addition, we determined the transcriptomic response of the same resistant cultivar (Chinese Spring) upon infection with *Z. tritici* wildtype strain (3D7) and with a mutant line expressing the avirulent allele of *AvrStb6* (3D7+AvrStb6_1E4_) at 3 and 6 dpi. To visualize the overall changes in the host transcriptome upon infection or protein infiltration, we performed a multidimensional scaling (MDS) on the expression data of the 33,249 expressed genes. The MDS showed that samples clustered according to experiments (*Z. tritici* infection assay vs AvrStb6-infiltration experiment, first dimension) and to timepoints of the spray inoculation experiment (second dimension; Supplementary Figure S6A). When the MDS was performed for each experiment separately, the data clustered according to the timepoints and the treatments (Supplementary Figure S6B and C). The MDS analysis indicated that AvrStb6-triggered transcriptomic reprogramming changes over time.

**Figure 4.**
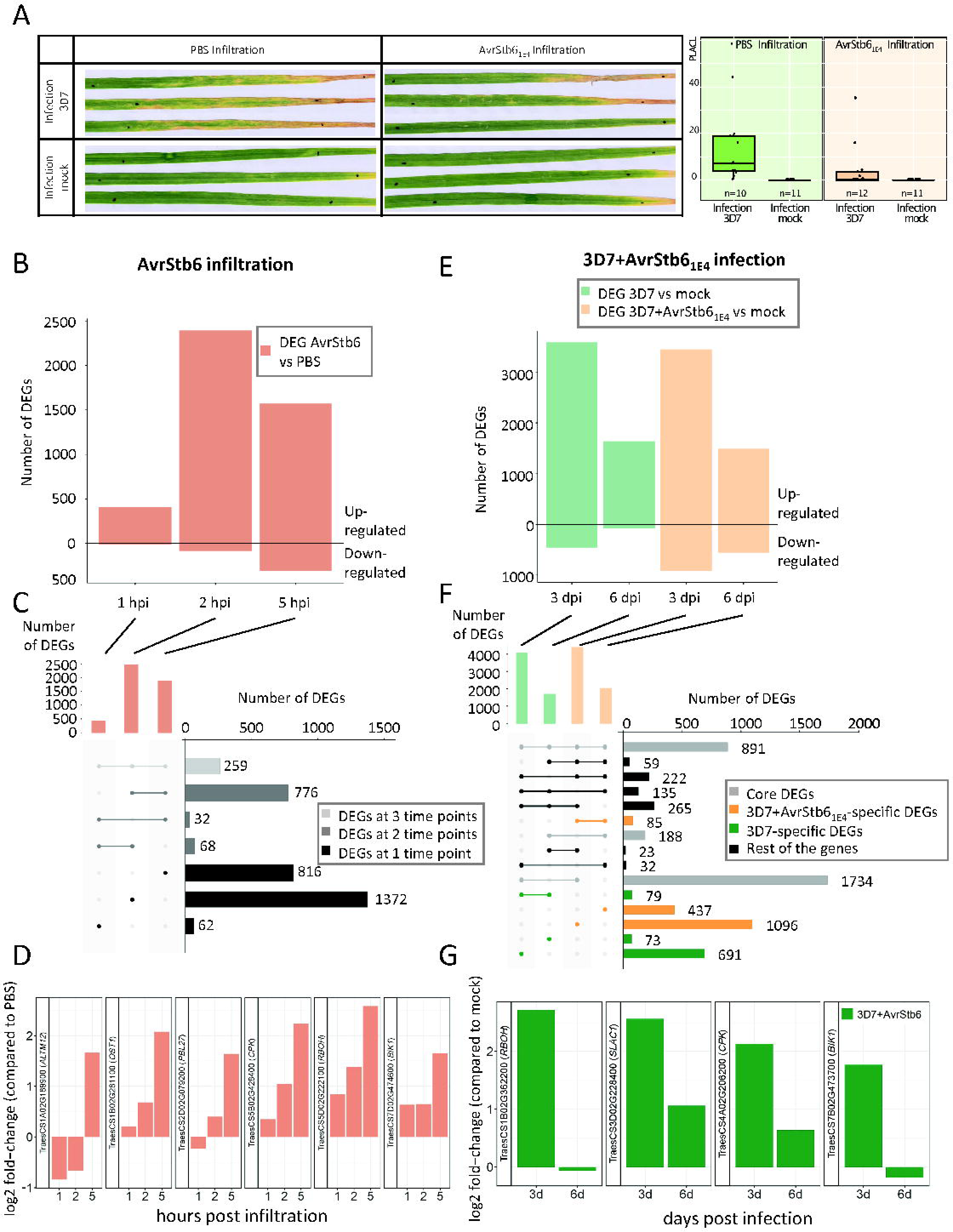
AvrStb6 induces an immune response in Stb6-expressing wheat plants. **A.** AvrStb6 produced in *Pichia pastoris* induces an immune response in the cultivar Chinese Spring and prevents the infection of *Zymoseptoria tritici* strain 3D7. AvrStb6 was infiltrated in Chinese Spring plants. Plants were subsequently treated with mock or with *Z. tritici* 3D7 spore suspension. Pictures were taken 11 days after infection. Virulence is estimated as percentages of leaf area covered by lesions (PLACL). **B.** Barplot showing the number of differentially expressed genes (DEGs) in wheat upon AvrStb6 infiltration compared to the PBS control at three different timepoints (1 hour post infiltration (hpi), 2hpi and 5 hpi). **C.** Intersection plot showing the overlap between the different sets of DEGs. Colors of bars in the right panel correspond to genes differentially expressed at the three timepoints (light grey), at two timepoints (dark grey) or at only one timepoint (black). **D.** Expression levels (log2 fold-change compared to PBS) of stomatal closure-related genes differentially up-regulated upon AvrStb6 infiltration at 5 hpi (log2 fold change >1.5; FDR < 0.01). Expression patterns of *ALMT12* (TraesCS1A02G189900), *OST1* (TraesCS1B02G281100), *PBL27* (TraesCS2D02G079200), 1 *CPK* (TraesCS5B02G428400), 1 *RBOH* (TraesCS5D02G222100), and 1 *BIK1* (TraesCS7D02G474600) homologs are shown. The expression values are at 1, 2 and 5 hours post infiltration of the purified AvrStb6. **E.** Barplot showing wheat DEGs upon infection with the virulent strain 3D7 and with the avirulent strain 3D7+AvrStb6_1E4_ compared to mock-treated plants at 3 or 6 days post infection (dpi). **F.** Intersection plots show the overlap between the different sets of DEGs. Colors of bars in the right panel correspond to three sets of genes: grey = core DEGs, orange = 3D7+AvrStb6_1E4_-specific DEGs and green = 3D7-specific DEGs. The rest of the genes are shown in black. **G.** Expression (log2 fold-change compared to mock) at 3 and 6 dpi of stomatal closure-related genes identified in “3D7+AvrStb6_1E4_-specific DEGs” set at 3 dpi. Expression patterns of 1 *RBOH* (TraesCS1B02G362200), *SLAC1* (TraesCS3D02G228400), 1 *CPK* (TraesCS4A02G206200), and 1 *BIK1* (TraesCS7B02G473700) homologs are shown. Genes shown are differentially expressed upon infection with the avirulent strain compared to the mock (log2 fold change >1.5; FDR < 0.01), but not with the virulent strain.

We first analyzed the transcriptomic response of wheat upon infiltration with the avirulent isoform of AvrStb6 protein. We performed differential gene expression analyses on the control plants (infiltrated with PBS) and AvrStb6-infiltrated plants at each timepoint (Supplementary Table S5). We identified 421, 2475, and 1883 differentially expressed genes (DEGs) at 1, 2, and 5 hpi, respectively, resulting in 3385 genes that are differentially expressed (DE) upon AvrStb6 infiltration at least at one timepoint (hereafter called “DEGs in infiltration”, Figure 4B and Supplementary Table S4). Out of these 3385 genes, 259 are differentially expressed at all the three timepoints and 876 at two timepoints. The results suggest that AvrStb6 triggered distinct waves of differentially regulated genes, most of them being upregulated (Figure 4C). DEGs after AvrStb6 infiltration were enriched in “calcium ion transmembrane transport”, “oxylipin biosynthetic process” Gene Ontology (GO) terms and other GO terms linked to biotic stress responses, photosynthesis, and cell wall-related terms (Supplementary Figure S7 and Table S6, “DEGs in infiltration”), suggesting that cell wall remodeling and immune response induction are key parts of the host response against AvrStb6. Remarkably, we identified several up-regulated genes involved in stomatal closure, i.e., seven genes encoding for putative *Respiratory Burst Oxidases* (*RBOH*), an *Open Stomata1* (*OST1*, protein kinase mediating the regulation of stomatal aperture/closure), 1 *Serine/threonine-protein kinase PBL27* , 4 *Calcium-Dependent Protein Kinase* (*CPK*), 1 *Aluminum-Activated, Malate Transporter 12* (*ALMT12*) and a Serine/threonine-protein kinase *Botrytis-induced Kinase1* (*BIK1*) homologs (Mustilli et al. 2002; Kwak et al. 2003; Geiger et al. 2010; Meyer et al. 2010; Sasaki et al. 2010; Brandt et al. 2012; Scherzer et al. 2012; Kadota et al. 2014; Zheng et al. 2018; Liu et al. 2019) (Figure 4D; Supplementary Table S7). These results suggest that wheat response to AvrStb6 infiltration may trigger Reactive Oxygen Species (ROS) accumulation and stomatal closure.

Second, we analyzed the transcriptome of wheat plants infected with the virulent strain 3D7 and the avirulent strain 3D7+AvrStb6_1E4_. In total, 4049 and 1713 genes were differentially expressed upon infection with the virulent strain 3D7 compared to the control treatment (mock) at 3 and 6 dpi, respectively (Figure 4E; Supplementary Table S5). In plants infected with the avirulent strain expressing AvrStb6 (3D7+AvrStb6_1E4_), 4375 and 2049 genes were differentially expressed from the control at 3 and 6 dpi, respectively (Figure 4E, and Supplementary Tables S4 and S5). We identified 3 distinct sets of differentially expressed genes: i) the “core DEGs’’ consisted of 2813 genes differentially expressed upon infection with both strains compared to the control at each timepoint (grey in Figure 4F); ii) the “3D7-specific DEGs’’ consisted of 843 genes differentially expressed exclusively upon infection with the virulent strain (green in Figure 4F); and iii) the “3D7+AvrStb6_1E4_-specific DEGs’’ were differentially expressed only upon infection with the AvrStb6-expressing strain, but not with the wild-type strain regardless of the timepoint (1618 genes, orange in Figure 4F). We performed a GO enrichment analysis in these three sets of genes to get insights into the function of the DEGs. We observed enrichment in core DEGs in GO terms linked to stress response and plant cell wall. We observed an enrichment in the “calcium ion transmembrane transport”, “oxylipin biosynthetic process”, “cellulose synthase” and “plant cell wall biogenesis” GO terms and in GO terms linked to photosynthesis only in the 3D7+AvrStb6_1E4_-specific DEGs (Supplementary Figures S7-S9). Within the 3D7+AvrStb6_1E4_-specific DEGs, up-regulated genes involved in stomatal closure regulation, including *Slow Anion Channel-associated1* (*SLAC1*), 2 *RBOH,* and 3 *CPKs* homologs were identified (Kwak et al. 2003; Negi et al. 2008; Vahisalu et al. 2008; Geiger et al. 2010; Brandt et al. 2012; Scherzer et al. 2012). In accordance with the previous findings with AvrStb6 protein, we conclude that recognition of the avirulent strain induces a transcriptomic reprograming involved in stomatal closure (Figure 4G; Supplementary Table S7).

Finally, the specific response to AvrStb6 was determined by combining the protein infiltration and infection results. Around 53% of the DEGs upon infiltration with the avirulent version of AvrStb6 protein were also differentially expressed in the presence of *Z. tritici* (1798 out of 3385 genes for the case of the avirulent strain, Figure 5A, B), highlighting that the direct infiltration of one single avirulence factor highly mimics the infection response in wheat. To determine the response triggered specifically upon AvrStb6 recognition, we identified 247 genes that were differentially expressed in the presence of AvrStb6 both when the avirulent version of AvrStb6 protein was applied (“DEGs in infiltration”) and when the plants were infected with the AvrStb6-expressing strain, but not with the virulent strain. These 247 genes were referred to as “AvrStb6 response” (Figure 5A, B). Of these, 103 and 76 genes were up-regulated and down-regulated, respectively, upon infection with the avirulent strain at 3 dpi. We checked the Uniprot annotation and blasted the protein sequences of all those 247 genes against the NCBI conserved domain database (Supplementary Table S8). Among the up-regulated genes upon infection by 3D7+AvrStb6 strain at 3 dpi that were also up-regulated upon AvrStb6 infiltration (Supplementary Table S9 and S10), we identified genes encoding for an MLO (Mildew resistance locus o)-like protein, for 1 chitinase domain-containing protein, for 2 NOD-like receptors (NLRs), for 1 putative receptor-like proteins (RLP) and for 24 putative receptor-like kinases (RLKs). Among the RLKs, 2 harbored a lectin-like domain, 17 a leucine-rich repeat (LRR) domain, 1 a jacalin-like protein kinase and 4 a wall-associated kinase (WAK) domain (Supplementary Table S9, highlighted in light green). Expression levels of some of these genes are shown in Figure 5C (Supplementary Table S10; highlighted in dark green). The previously cloned resistance genes *Stb6*, *Stb15* and *Stb16q* were up-regulated upon AvrStb6 treatment, and upon the infection by the avirulent strain and the virulent strain (Supplementary Table S4, in red, green and blue, respectively). Therefore, although these well-characterized resistance genes are up-regulated upon AvrStb6 recognition (Set_DEG_INF_all), they were not considered to be part of the AvrStb6-specific response. We additionally identified several up-regulated genes upon AvrStb6 infiltration and infection with the avirulent strain encoding proteins involved in cell wall modification, including 1 glycosyltransferase, 1 pectin esterase, 2 glucan endo-1,3-beta-D-glucosidases, and an alpha-L-fucosidase (Supplementary Table S9, highlighted in orange). Moreover, we identify some putative genes encoding proteins playing a role in stomatal movement, including two CBL-interacting protein kinase (CIPKs), a sugar transporter protein (STP), and a plant glutamate receptor (GLR) homolog (Kong et al. 2016; Förster et al. 2019; Flütsch et al. 2020) (Figure 6B; Supplementary Table S9, highlighted in blue). These results suggest that the specific response triggered upon AvrStb6 recognition involves a transcriptomic reprogramming characterized by the induction of defense-, stomatal closure-, and cell wall-related genes.

**Figure 5.**
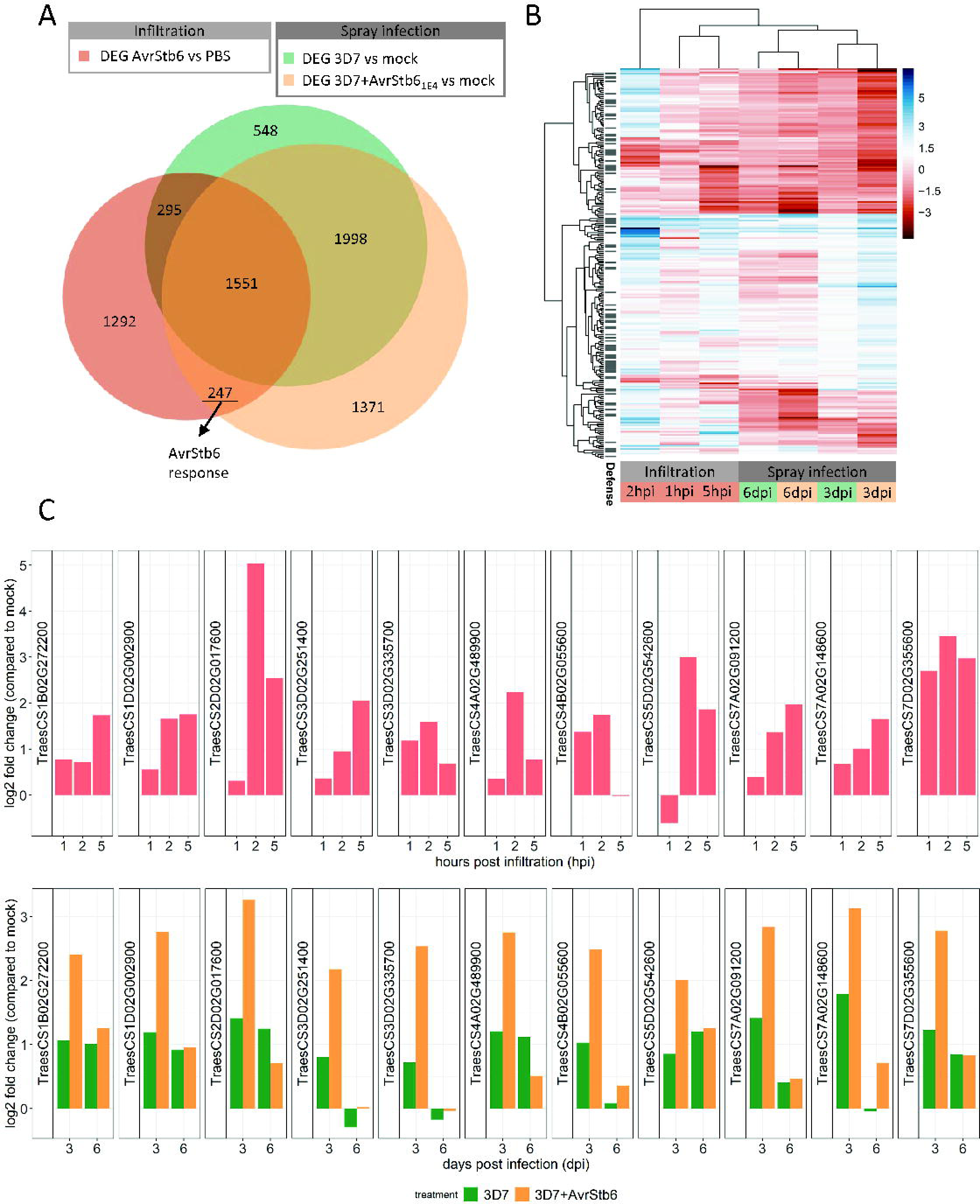
Defense-related genes contribute to AvrStb6-specific response in wheat. **A.** Venn diagram representing the overlap between the differentially expressed genes identified in the different pairwise comparisons: infiltration of AvrStb6 compared to control (PBS), infection with 3D7 compared to the mock treatment, and infection with the avirulent strain 3D7+AvrStb6_1E4_ compared to the mock treatment. The highlighted 247 genes correspond to the genes responding specifically to the presence of AvrStb6. **B.** Heatmap of the 247 genes differentially expressed in the presence of AvrStb6 in both the spray infection and infiltration experiments. The log2 fold change is represented for each pairwise comparison. In the left column, black bars indicate the genes containing protein domains linked to defense responses. **C.** Expression pattern of an *MLO* (*TraesCS1B02G272200*) and 10 *RLKs* within the 247 genes (differentially expressed upon infiltration with AvrStb6 protein and infection with 3D7+AvrStb61E4, but not with 3D7). Top panel: log2 fold change expression upon infiltration of AvrStb6 at 1, 2, and 5 hours post-infiltration (hpi). Bottom panel: log2 fold change expression upon infection with the wildtype strain (3D7) and the avirulent strain (3D7+AvrStb6) compared to mock at 3 and 6 days post-infection (dpi).

**Figure 6.**
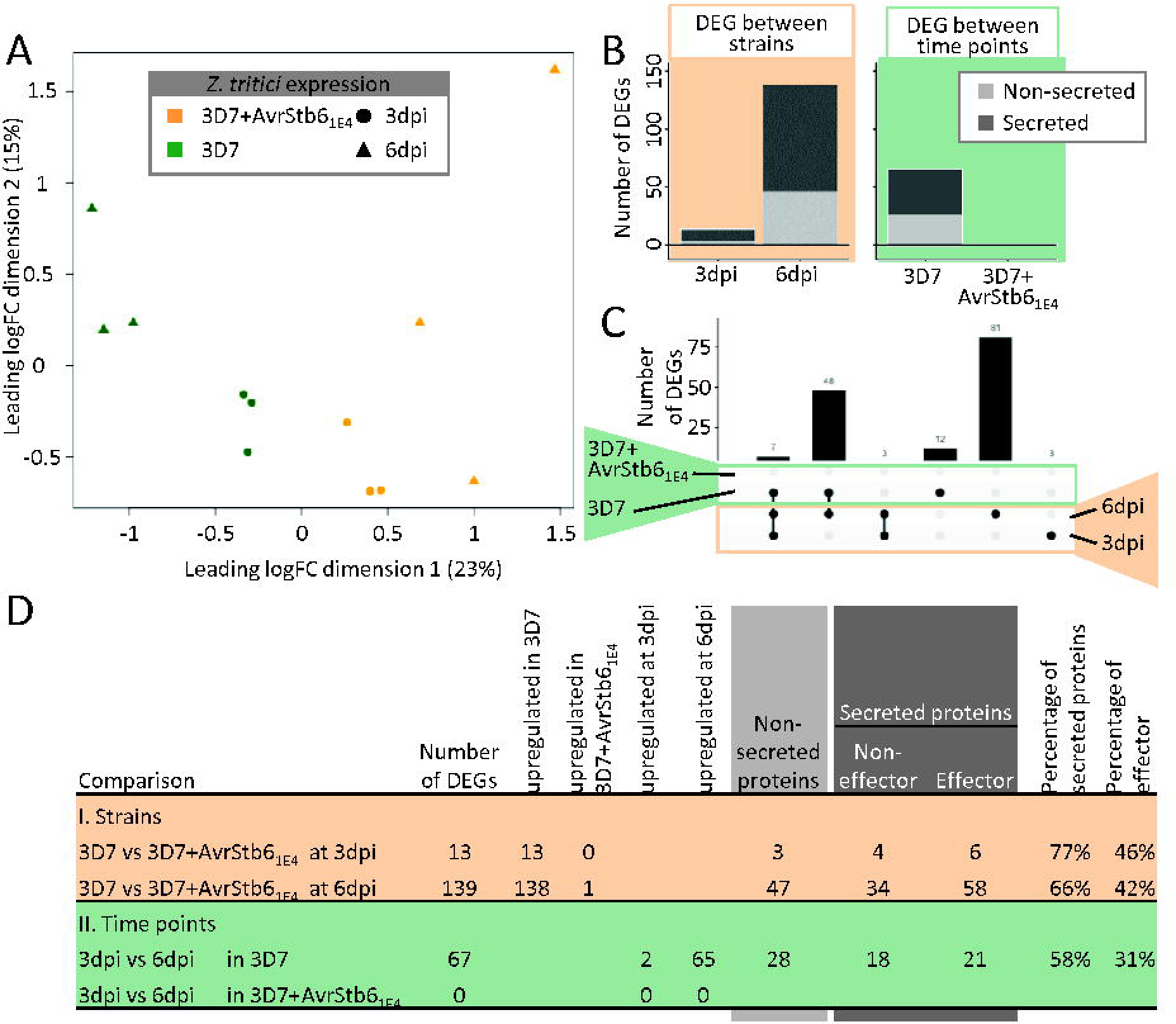
The avirulent AvrStb6 expressing strain is blocked at the early stages of the infection. **A.** Multidimensional scaling (MDS) plot of the expression data of *Zymoseptoria tritici* virulent (3D7) and avirulent (3D7+AvrStb6_1E4_) strains, during infection of the resistant cultivar Chinese Spring at 3 and 6 days post-infection (dpi). The different colors correspond to different treatments (green = 3D7 and orange = 3D7+AvrStb6_1E4_), representing the two timepoints with different shapes. **B.** Barplots showing the number of differentially expressed genes (DEGs) between strains (light orange) and between timepoints (light green). Light grey represents genes encoding not-secreted proteins, and dark grey genes encoding secreted proteins. **C.** Intersection plots showing the overlap between the different sets of DEGs (Green: between time points for each strain; Orange: between strains at each time point). **D.** Summary of the DEGs identified in the different comparisons and the percentage of secreted proteins and effector genes.

### Transcriptomic pattern of Z. tritici avirulent strains

We finally analyzed the transcriptomic pattern of virulent and avirulent *Z. tritici* strains during infection and showed that data clustered based on timepoint and genotype (Figure 6A). Remarkably, no major changes were identified in the transcriptome of the avirulent strain 3D7+AvrStb6_1E4_ between the two timepoints analyzed (Figure 6B and C and Supplementary Table S11). Thirteen genes were differentially expressed between 3D7 and 3D7+AvrStb6_1E4_ at 3 dpi and 139 at 6 dpi (Figure 6B). By comparing the lists of DEGs, we noticed that most of the genes (82%) that were differentially expressed between the two timepoints in 3D7 were also differentially expressed between the two strains at 6 dpi, suggesting that both strains have a similar transcriptomic profile at 3 dpi, but at later stages 3D7 transcriptomic profile changes, consistent with its successful host colonization (Figure 6C). We then analyzed the sequences of all the DEGs in *Z. tritici* for putative effector genes. Around 60% of the differentially expressed genes between both isolates were secreted, and more than 40% were putative effector genes (Figure 6D). The results indicate that the immune response triggered in wheat plants upon AvrStb6 recognition does not induce detectable changes in the transcriptome of *Z. tritici*.

## DISCUSSION

Resistance towards fungal pathogens features the specific recognition of avirulence factors, which prevents the progression of the infection. However, the mechanisms involved in arresting the growth of the avirulent strain remain largely unknown. In this work, we provide a better understanding of the intricate mechanisms that are involved in strain-specific resistance against the major wheat pathogen *Z. tritici*. We demonstrated that an accumulation of AvrStb6 effector occurs during penetration of the pathogen through the stomata. At this stage, the pathogen progression is arrested in resistant wheat cultivars harbouring Stb6. We propose that AvrStb6 triggers a rapid induction of wheat defense, cell wall-related genes, and genes involved in stomatal closure, contributing to preventing host colonization of the avirulent strain and its penetration through the stomata.

*Z. tritici* has an extended asymptomatic phase considered critical for the outcome of the infection (Sánchez-Vallet et al. 2015; Steinberg 2015). During this phase, *Z. tritici* spores germinate, hyphae grow epiphytically on the leaf surface, penetrate through the stomata or wounds, and grow in the apoplastic space (Kema et al. 1996; Duncan and Howard 2000; Fones et al. 2017; Haueisen et al. 2019; Battache et al. 2022). The cytological analysis carried out in this work enabled us to identify the stomata as key for the infection process, not only because it is the most frequent gate for fungal penetration (Kema et al. 1996; Duncan and Howard 2000; Fones et al. 2017), but also since it is where fungal cells accumulate AvrStb6, a critical avirulence factor, and where the growth of avirulent strains is arrested. Indeed, we observed that epiphytic hyphae produce very low levels of AvrStb6, while cells attempting to penetrate the stomata accumulate AvrStb6. Once the virulent strains entered the substomatal cavity and colonized the apoplast, AvrStb6 levels remained high. These results are in accordance with the reported gene expression pattern of *AvrStb6* (Meile et al. 2020) and are similar to the one described for another avirulence gene, *Avr3D1* (Meile et al. 2023). Previously published transcriptomic analyses showed that both effector genes (*AvrStb6* and *Avr3D1*) were induced at the onset of the necrotrophic phase (Rudd et al. 2015; Palma-Guerrero et al. 2017; Zhong et al. 2017; Meile et al. 2018). However, literature (Meile et al. 2020, 2023) and our microscopic analysis indicate that these two effectors are induced much earlier during the infection. We hypothesize that other effector genes with similar expression patterns (Rudd et al. 2015; Palma-Guerrero et al. 2017; Haueisen et al. 2019) could also be activated during penetration and remain relatively high during growth in the apoplast.

Taken together, the restricted expression pattern of *AvrStb6*, the observed AvrStb6 accumulation in fungal cells close to stomata, and the blockage of AvrStb6-expressing strains at the level of the stomata indicate that AvrStb6 is recognized during stomatal penetration. Plants regulate stomatal opening upon a pathogen attack to prevent penetration by those pathogens that use natural openings, indicating that pathogen detection through stomata is a general defensive strategy (Melotto et al. 2017; Wu and Liu 2022). Noticeably, it was recently reported that AvrStb16 and Avr3D1 recognition also block avirulent isolates during wheat invasion through stomata and that wheat stomata were closed following recognition of *Z. tritici* strains expressing AvrStb16 or AvrStb6 (Saintenac et al. 2021; Battache et al. 2022; Ghiasi Noei et al. 2022; Meile et al. 2023). We propose that AvrStb6 recognition occurs at the stomata and that resistance proteins recognizing Avr3D1, AvrStb6 and AvrStb16 are expressed in both guard cells and subsidiary cells. We identified several genes involved in stomatal closure that were upregulated upon AvrStb6 treatment and upon infection with the avirulent strain, including seven homologs of RBOH, 1 homolog of BIK1, 2 homologs of PBL27, 3 homologs of CPK, 1 homolog of OST1, 1 homolog of ALMT12, and 1 homolog of SLAC1 (Mustilli et al. 2002; Kwak et al. 2003; Negi et al. 2008; Vahisalu et al. 2008; Geiger et al. 2010; Meyer et al. 2010; Sasaki et al. 2010; Brandt et al. 2012; Scherzer et al. 2012; Kadota et al. 2014; Zheng et al. 2018; Liu et al. 2019). These results suggest that stomatal closure triggered by the avirulent strain and AvrStb6 treatment could be due to recognition by transmembrane receptors that, in turn, could activate NADPH oxidases, protein kinases, and anion channels to initiate a ROS burst and promote stomatal closure.

Our transcriptomic analysis, which combines the treatment with AvrStb6 protein and the infection with isogenic *Z. tritici* lines with and without *AvrStb6*, revealed the specific transcriptomic response upon recognition of this avirulence factor. Among the genes differentially expressed upon AvrStb6 infiltration and infection with the avirulent strain, cell wall-related GO terms were enriched. We hypothesize that changes in the cell wall triggered by recognition of the avirulence factor result in arresting fungal growth, as shown in other pathosystems (Menna et al. 2021; Molina et al. 2021). However, we cannot discard that AvrStb6 function involves targeting the cell wall independently of the immune response triggered upon its recognition. We identified several resistance-related genes up-regulated upon infection by the avirulent strain and treatment with the avirulence factor. In particular, we identified 24 RLKs and 1 RLPs which are known to be crucial for plant immunity (Tang et al. 2017). Remarkably, the resistance genes *Stb6*, *Stb16q,* and *Stb15* were up-regulated upon AvrStb6 infiltration and infection with the virulent and avirulent strains of *Z. tritici*, suggesting that the resistance genes are induced during wheat infection (Bernasconi et al. 2023) but also upon recognition of AvrStb6.

The dual transcriptomic approach simultaneously determined the response triggered in the plant and its effect on the pathogen. Our results showed that the virulent and avirulent strains harbor different acclimation strategies to the host at the transcriptomic level. At an early timepoint (3 dpi), both the virulent and avirulent hyphae mostly grow epiphytically. Accordingly, at this timepoint, only 13 genes were differentially expressed between the virulent and the avirulent strain, of which 6 were predicted to be effectors. At a later timepoint (6 dpi), the virulent strain colonizes the apoplast. However, the avirulent strain remains blocked on the leaf surface. This translates into a new set of genes induced in 3D7 that remain unchanged in the avirulent strain. Among this new set of genes, 58 are predicted to encode putative effectors and are probably involved in penetration or apoplast colonization of virulent strains. We expected an induction of stress-related genes in the avirulent strain, but we were not able to detect a transcriptomic reprogramming. We hypothesize that this lack of transcriptomic response is due to the fact that the defense response only impacts upon specific fungal cells (e.g. cells attempting to penetrate), and that this response was not detected in the whole leaf transcriptomic approach undertaken in this study. Alternatively, the observed phenotype might indicate that *Z. tritici* does not undergo a strong stress response upon recognition.

## Conclusions

This work describes how gene-for-gene resistance acts in wheat against the major pathogen *Z. tritici*. Our results shed light on the resistance response triggered upon pathogen recognition and suggest that cell wall modifications might play a key role. Furthermore, we provided evidence for the capacity of fungal strains to sense the apoplast and/or the stomata to produce effector proteins. Finally, our data demonstrate that recognition of avirulent strains prevents their penetration through the stomata and, subsequently, apoplast colonization.

## MATERIALS AND METHODS

### Plant and Fungal material

Wheat seeds (*Triticum aestivum*) of cultivars Chinese Spring and Runal were provided by DSP Ltd. (Delley, Switzerland). Seeds of the 2 Bobwhite near-isogenic lines (Bobwhite, Bobwhite + Stb6 T2-19-2, and Bobwhite + Stb6 T2-41-2) (Saintenac et al. 2018) were kindly provided by Kostya Kanyuka and bulk-amplified in our greenhouse facility (16h photoperiod, 70% humidity, 18°C during the day and 15°C at night). For all experiments described in this study, wheat seeds were grown in square pots (11 x 11 x 12 cm; about 11 plants per pot) containing the peat substrate Jiffy GO PP7 (Jiffy Products International, Moerdijk, the Netherlands). Plants were grown on a 16h photoperiod at 18°C (day) and 15°C (night) with 80% humidity and 30 klx.

All *Zymoseptoria tritici* strains used in this study were derived from the Swiss isolates ST99CH_3D7 and ST99CH_1E4 (abbreviated as 3D7 and 1E4, respectively) (Linde et al. 2002; Zhan et al. 2002). The reporter lines 3D7-GFP, 3D7-mCherry, 1E4-GFP, and 3D7-mTurquoise + p*AvrStb6*_1E4_::*His1-mCherry* were previously described (Meile et al. 2020; Barrett et al. 2021). Strains expressing AvrStb6 variants (AvrStb6_1E4_, AvrStb6_1E4_-GFP, and AvrStb6_3D7_-GFP) were engineered specifically for this study as described below. *Z. tritici* blastospores were grown in 50ml liquid cultures with yeast-sucrose broth medium (YSB; 10 g L^-1^ yeast extract, 10 g L^-1^ sucrose) or on solid yeast malt agar medium (YMA; 4 g L^-1^ yeast extract, 4 g L^-1^ malt extract, 4 g L^-1^ sucrose, 12 g L^-1^ agar), in the dark, at 18°C. Culture media were supplemented with 50 µg mL^-1^ kanamycin sulfate. Liquid cultures were grown with agitation (250 rpm). *Escherichia coli* and *Agrobacterium tumefaciens* were grown in Luria-Bertani medium (LB; 5 g L^-1^ yeast extract, 10 g L^-1^ tryptone, 10 g L^-1^ NaCl, pH7.5) supplemented with 50 µg mL^-1^ kanamycin sulfate or with a combination of 50 μg mL^−1^ kanamycin sulfate, 100 μg mL^−1^ carbenicillin and 50 μg mL^−1^ rifampicin, respectively. LB was supplemented with 10 g L^-1^ agar when used as solid media. *E. coli* and *A. tumefaciens* were grown in the dark at 37°C and 28°C, respectively. All strains are listed is Supplementary Table S1

### Establishment of Z. tritici AvrStb6 reporter lines

The reporter line ectopically expressing *AvrStb6* from the avirulent strain 1E4 (*AvrStb6*_1E4_), under the control of its native promoter and terminator regions (1010 and 1009 bp upstream and downstream of *AvrStb6*_1E4_ coding sequence, respectively), was obtained by transforming 3D7-GFP (Barrett et al. 2021) with the vector pES1_Avr1E4 previously described (Zhong et al. 2017). To obtain the strains expressing AvrStb6 fused to the Green Fluorescent Protein (GFP) reporter (AvrStb6_1E4_-GFP and AvrStb6_3D7_-GFP), 2 plasmids containing the *AvrStb6* promoter (1009 bp), the open reading frame (ORF) without the stop codon, and the terminator region (1010 bp) of 1E4 and 3D7, respectively, were engineered. 3D7 and 1E4 sequences were amplified from pES1_Avr1E4 (Zhong et al. 2017) and 3D7 genomic DNA, respectively. The *Z. tritici* codon-optimized GFP was amplified from pCeGFP (Kilaru et al. 2015) using NEB Phusion polymerase (New England Biolabs) and the primers listed in Supporting Information Supplementary Table S2. A linker (GGSGGGSG) was integrated between *AvrStb6* and *GFP*. PCR products were purified using the NucleoSpin Gel and PCR Cleanl1Jup kit (MACHEREY-NAGEL) and introduced into linearized (with KpnI) pCGEN (Motteram et al. 2011) using the In-Fusion HD Cloning Kit (Takara Bio). The resulting vectors, JA031_pCGEN_p1E4-1E4-GFP-t1E4 and JA035_pCGEN_p3D7-3D7-GFP-t3D7 (Genebank files available in doi: 10.3929/ethz-b-000641399), were used to transform chemically competent *E. coli* HST08 (Stellar competent cells, Takara Bio). Verified plasmids using Sanger sequencing were electroporated into *A. tumefaciens* AGL1 cells and were used to transform the *Z. tritici* reporter line 3D7-mCherry (Barrett et al. 2021). *A. tumefaciens* -mediated transformation was performed as described previously (Meile et al. 2018). Insertions were confirmed by Sanger sequencing. Transgene copy numbers were assessed by quantitative PCR (qPCR), using primers specific for the resistance marker gene (hygromycin B phosphotransferase gene, *hph*, for the untagged *AvrStb6* transformant and neomycin resistance gene, *neo*, for *AvrStb6-GFP* transformants) and the reference gene *TFIIIC1* (*Mycgr3G110539*).

Quantitative PCRs were performed on Roche LightCycler480 following the HOT FIREPol EvaGreen qPCR recommendations (primers listed in Supplementary Table S2). Only transformants with a single transgene insertion were selected for further analysis. We confirmed that the fused proteins were functional since the lines expressing *AvrStb6*_1E4_*-GFP* were avirulent in the cultivar Chinese Spring, but not in the susceptible cultivar Drifter (Supplementary Figure S1).

### Infection assays

Plants were sprayed with a suspension of 10^6^ spores mL^−1^ in 0.1% Tween 20 until run-off following the procedure described (Meile et al. 2018). For mock treatments, plants were sprayed with a 0.1% Tween 20 solution. For symptom quantification, second or third leaves were mounted on paper sheets, scanned with a flatbed scanner (CanoScan LiDE 220), and analyzed following the method described in (Zenkl et al. 2023). Data analysis and plotting were performed using RStudio 2023.06.1+524 (RStudio 2020).

### AvrStb6 protein secretion assay

*Z. tritici* strains expressing either *AvrStb6*_1E4_*-GFP* or *AvrStb6*_3D7_*-GFP* were grown in liquid medium, as mentioned in the fungal material section. Seven days after inoculation, the axenic cultures were filtered through 2 layers of sterile cheesecloth and pelleted (3273 g, 15 min, 4°C). Supernatants (SN) were filtered using 0.2-µm filters, lyophilized and resuspended in 1.5 ml Tris-buffered saline (TBS; 25 mM Tris, 150 mM NaCl, pH 7.5) before aliquoting (60 µL) and stored at -80°C until further experiments (“Filtered SN”). Pellets were washed four times with sterile deionized water and resuspended in 10 ml breaking buffer (BB; 50 mM sodium phosphate, 1 mM phenylmethylsulfonyl fluoride, 1 mM EDTA, 5% glycerol, pH 7.4). Cells were pelleted again (10 min, 3000 g, 4°C) and resuspended in BB to reach an OD_600_ of 100. An equal volume of glass beads was added to the samples before proceeding with 8 cycles of 30 seconds of vortex homogenization followed by 30 seconds on ice. Samples were then centrifuged at 12000 g for 10 min, and the collected supernatant corresponded to the “Washed pellets” without urea samples, and the remaining pellet was treated with urea. Pellets were resuspended in 10 mL fresh BB supplemented with 6 M urea, and the same procedure was applied (starting from the vortex/ice cycles) to obtain the “Washed pellets” with urea samples. Immunoprecipitation of the samples was performed following the GFP-Trap_MA protocol (Chromotek) using 6 µL of beads per sample and equilibrating the samples with 1.5 volumes of GFP-Trap_MA wash buffer. Elution was done in 2x Laemmli Buffer (125 mM Tris Base, 4% SDS, 20% glycerol, 10% 2-mercapto-ethanol, 2mg mL^-1^ Bromophenol blue).

### Protein production in Pichia pastoris

The recombinant protein AvrStb6_1E4_, tagged with a C-terminal c-myc epitope and a polyhistidine (6xHis) tag was purified using the *Pichia pastoris* heterologous expression system. The *AvrStb6*_1E4_ coding sequence lacking its endogenous signal peptide was amplified from 1E4 cDNA and pPICZalphaB (Invitrogen) was linearized using an inverse PCR approach. Amplifications were performed using NEB Phusion polymerase with the primers listed in Supplementary Table S2, gel-purified with the NucleoSpin Gel and PCR Cleanl1Jup kit (MACHEREY-NAGEL) and assembled using the In-Fusion HD Cloning Kit (Takara Bio). After Sanger sequencing validation, the resulting vector (pPICZalphaB_AvrStb6_1E4; Genebank file available in supplementary files) was used for transformation of *Pichia pastoris* strain X-33, which was cultured according to the *Pichia* Expression Kit manual (K1710-01, Invitrogen).

*P. pastoris* preculture was prepared in 250 ml yeast extract peptone dextrose (YPD; 1% yeast extract, 2% peptone, 2% dextrose) media at 28°C, 250 rpm, until reaching an OD_600_ of between 2 and 6 (for 16-18 h). The preculture was pelleted at 3000 g for 5 min, the cells were resuspended in buffered glycerol-complex medium (BMMY; 1% yeast extract, 1% peptone, 100 mM potassium phosphate, pH 6, 1.34% yeast nitrogen base medium, 4×10^-5^ % biotin, 1% methanol) to a final OD_600_ of 1, divided in eight 250mL cultures (grown in 1L sterile Erlenmeyer) and grown for 29 h at 28°C and shaking at 265 rpm. Methanol was added to a final concentration of 0.5% after 24 h. The supernatant of the culture was centrifuged at 13689 g for 4 min at 4°C. For AvrStb6 purification, we followed the HisPur Ni-NTA resin batch protocol (Thermo Scientific). Ni-NTA-bound AvrStb6 was eluted in phosphate-buffered saline (PBS; 20 mM sodium phosphate, 300 mM sodium chloride pH 7.4) supplemented with 250 mM imidazole. Imidazole was eliminated from elution fractions by buffer exchange using PBS and illustra NAP-25 columns (GE Healthcare). The presence of purified protein in the elution fractions was confirmed by western blot analysis using anti-HA antibodies (western blot analysis details below). The concentration of the purified protein was estimated by measuring the 280-nm absorbance and accounting for the cysteine reductions. To evaluate the functionality of the purified AvrStb6_1E4_, we infiltrated 13 days post germination (dpg) Chinese Spring second leaves with either the purified AvrStb6 recombinant protein at a concentration of 176 mg mL^-1^ (23 leaves) or PBS (21 leaves). Subsequently, plants were spray-inoculated with 3D7 or with a mock treatment as described in the infection assay section. Second leaves were harvested 11 days after infection for symptom evaluation.

### SDS-PAGE, western blots, and Coomassie analysis

Protein samples were separated by SDS-PAGE and transferred to a polyvinylidene fluoride (PVDF) membrane (TurboMidi 0.2 µm PVDF membrane; BioRad) using the Trans-Blot Turbo system (Bio-Rad). Membranes were blocked at room temperature (RT) in blocking buffer (PBS, 0.01% Tween 20, 4% BSA) and incubated in a primary antibody solution anti-GFP-HRP (1/5000, MACS Miltenyi Biotec) and 1/5000 anti-His (11667475001, Roche) for GFP and His-Tag detection, respectively) followed by incubation with anti-mouse IgG (1/10000). SuperSignal West Pico Chemiluminescent Substrate kit (Thermo Scientific) was used for immunodetection. Blotted membranes were stained with Coomassie staining solution (10% acetic acid, 45% EtOH, 0.1% Brillant blue R, 20 min, RT). Image acquisition was performed on a ChemiDoc Imaging System (Bio-Rad).

### Confocal laser scanning microscopy

Second leaves of infected and mock-treated plants were harvested immediately before observation. The top 3 cm of each leaf were discarded, and the adaxial side of the adjacent section of approximately 2 cm was observed in 0.02% Tween 20 solution. Propidium iodide (PI) staining was performed by soaking the leaf sections in 0.02% Tween 20 solution amended with 10 µg mL^-1^ PI and applying vacuum. Images were acquired using an inverted Zeiss LSM 780 confocal microscope. Images of the *AvrStb6* promoter activity were acquired as described (Meile et al. 2020). All other images were acquired using two excitation sources, a diode-pumped solid-state laser (DPSSL; 561 nm) and an argon (488 nm) laser using the following detection settings: 494.95-535.07 nm for GFP signal, 623.51-641.26 nm for mCherry signal, 656.01-681.98 nm for the chloroplast autofluorescence, and 596.87-632.38 nm to detect PI signal. Images were processed using the Fiji platform of ImageJ (Schneider et al. 2012). Image processing included brightness and contrast adjustments, median filters (radius of 1 pixels), generation of maximum intensity z-projection, orthogonal projections, cropping, and addition of scale bars.

### Plant inoculation, tissue harvest, and RNA sequencing

Chinese Spring wheat plants (12 days post sowing) were treated with 3D7-GFP, 3D7-GFP expressing *AvrStb6*_1E4_ (3D7-GFP+*AvrStb6*_1E4_) or mock solution (0.1% (v/v) Tween 20 solution), as described for the infection assays. Samples were collected at 3 and 6 days post infection (dpi). For each treatment, the top 3 cm from the leaf tips were discarded, three second leaves (from independent plants) were pooled and flash-frozen in liquid N_2_. Concurrently, 13-day old Chinese Spring leaves were infiltrated with 176 mg mL^-1^ of AvrStb6 purified protein (see Protein production in *P. pastoris*) or with PBS buffer. Infiltrated areas were marked and collected at 1, 2 and 5 hours after infiltration and flash-frozen in liquid N_2_. Samples were homogenized with zirconium beads (1.4 mm diameter) using a Bead Ruptor equipped with a cooling unit (Omni International, Kennesaw, GA, USA). Total RNA isolation was performed using GENEzol reagent (Geneaid Biotech, Taipei, Taiwan) following the manufacturer’s instructions. DNA contamination was removed using the on-column DNase treatment of the RNase-Free DNase Set (Qiagen GmbH, Hilden, Germany). Concentration and quality were measured by Qubit fluorometer (Life Technologies) and Tapestation (Agilent). Library preparations and sequencing were performed at the Functional Genomics Center Zurich (http://www.fgcz.ch/). Ribosomal RNA was depleted by poly-A enrichment and DNA libraries were sequenced. The library preparation was performed with Illumina’s TruSeq Stranded mRNA kit using 500 ng of total RNA as input amount. The sequencing of the resulting libraries was performed on Illumina NovaSeq 6000 in single read 100 bp mode.

### RNA-seq analysis

Raw Illumina reads were pseudo-aligned to the wheat transcriptome (release 41 on https://plants.ensembl.org/, accessed November 2018) and to the *Z. tritici* transcriptome (Accession number PRJNA290690 on https://www.ncbi.nlm.nih.gov/, accessed November 2018) using Kallisto v0.44.00 (Bray et al. 2016; Supplementary Table S3). Read counts were then imported into R using the R-package tximport (Soneson et al. 2015; RStudio 2020). Normalization for RNA composition was performed with the edgeR package (Robinson et al. 2010) using the function calcNormFactors, and dispersion was estimated using the following functions: estimateGLMCommonDisp, estimateGLMTrendedDisp, and estimateGLMTagwiseDisp. For both wheat and *Z. tritici,* a gene was considered expressed if the counts per million (cpm) were >5 in at least 3 out of the 18 replicates, as described (Poretti et al. 2021).

Differential gene expression analyses were performed using both edgeR (Robinson et al. 2010) and DESeq2 (Love et al. 2014). For edgeR, the pipeline described (Praz et al. 2017) was followed. A negative binomial generalized log-linear model has been fitted to the read count for each gene using the glmFit function, and we tested for differential expression with the glmLRT function with different contrasts for pairwise comparisons. In DESeq2, standard parameters were used. Genes with a log_2_FC > |1.5| and an adjusted p-value (FDR) < 0.01 in both approaches (DESeq2 and edgeR) were included in the final set of differentially expressed genes (DEGs; Supplementary Table S4). Various pairwise comparisons were performed between infiltrated or infected samples against the corresponding control. The different pairwise comparisons are described in Supplementary Table S5. All raw sequence data generated in this study have been deposited in the NCBI Sequence Read Archive under accession number GSE250605.

### Lists of the genes of interest and their functional annotation

To functionally annotate the genes identified as DE in wheat, GO annotations were retrieved from Ensembl using the R package BiomaRt (Durinck et al. 2009), and GO enrichment analyses were performed for the “DEGs in infiltration”, the “Core DEGs”, the “3D7+AvrStb6_1E4_-specific DEGs’’ and the “3D7-specific DEGs’’ using topGO (Alexa and Rahnenfuhrer 2022). GO terms identified as enriched in any of the lists of genes of interest were manually classified into different categories according to their biological functions (Supplementary Table S6). To functionally annotate the “AvrStb6 specific genes’’, we used Uniprot (https://www.uniprot.org/) and performed blast searches against the conserved domain database (CDD, https://www.ncbi.nlm.nih.gov/cdd/) using standard parameters (Supplementary Tables S8, S9 and S10). Stomata related genes and NADPH oxidase genes were identified by BLAST using the Arabidopsis sequence as a query. To identify secreted proteins and effector candidates in the DEGs in *Z. tritici*, the protein sequences of all the genes identified as DE in each comparison have been analyzed using SignalP version 6.0 (Teufel et al. 2022) and EffectorP version 3.0 (Sperschneider and Dodds 2022).

## ACKNOWLEDGEMENTS

We thank Sreedhar Kilaru and Gero Steinberg for providing pCmCherry, pCZtGFP, and strains 3D7 and 1E4 expressing GFP; and Kostya Kanyuka for providing Bobwhite and Bobwhite+Stb6. Seeds of cultivars Runal and Chinese Spring were kindly purchased from DSP Ltd. (Delley, Switzerland). We thank Jason Rudd for providing pCGEN. Confocal laser scanning microscopy experiments were supported by the Scientific Center for Optical and Electron Microscopy (ScopeM), ETH Zurich. RNA extraction and RNAs-eq analysis were performed in collaboration with the Genetic Diversity Center (GDC), ETH Zurich. RNA sequencing was conducted in the Functional Genomics Center Zurich (FGCZ).

This work was supported by the Ministry of Science and Innovation (Grant PID2019-108693RA-I00 to ASV). ASV was the recipient of the RYC2018-025530-I grant from the Spanish Ministry of Science, Innovation and Universities. CP was a recipient of the Early Postdoc. Mobility-Grant, Nr. P2ZHP3_195287, from the Swiss National Science Foundation. ADF obtained the “*Ayudas María Zambrano para la atracción de talento internacional*“ Postdoctoral Fellowship from the Ministry of Universities of the Spanish government and the European Union Next Generation EU. CCL was financially supported by the ‘Severo Ochoa (SO) Programme for Centres of Excellence in R&D’ from the Agencia Estatal de Investigación of Spain (grant CEX2020-000999-S (2022-2025) to the CBGP).

## Figure legends

**Supplementary Figure S1. Virulence assays of the mutant lines obtained in this work. A**. Virulence estimated as percentage of leaf area covered by lesions (PLACL) of 3D7-mCherry lines expressing AvrStb6 alleles from 3D7 or 1E4 (*AvrStb6*_3D7_ or *AvrStb6*_1E4_) fused to GFP on Chinese Spring and Drifter wheat cultivars. 1E4, 3D7-mCherry, and mock treatments are shown as controls. **B** Virulence (estimated as PLACL) of 3D7-GFP expressing the avirulent *AvrStb6* allele from 1E4 (3D7-GFP_AvrStb6_1E4_ lines A1 and line A2) on wheat plants cultivars Chinese Spring and Drifter. 1E4, 3D7-GFP, and mock were included as controls. **C** Virulence assay of 1E4 strain on the wild-type Bobwhite cultivar and two independent lines (T2_19-2 and T2_41-4) of Bobwhite expressing Stb6. Expected compatible and incompatible interactions are highlighted in brown and green, respectively. Dark and light colors indicate the transformant lines and the background lines, respectively. The number of leaves analyzed is indicated for each condition (n number).

**Supplementary Figure S2. Rare events of penetration of** Z. tritici **in an incompatible interaction.** Confocal microscopy images of Chinese Spring sprayed with the *Z. tritici* cytosolic reporter line 1E4 (1E4-GFP). **A**. At 10 days post-infection (dpi), rare events of 1E4 having penetrated the leaf through stomata can be observed. Apoplastic hyphae remain in close proximity to the penetration site. **B.** At 14 dpi fungal progression in the apoplast remains very limited. Images A and B are maximum projections, A’ and B’ are orthogonal views. Dashed lines indicate the location of the orthogonal view on the corresponding image. Images are overlays of GFP signal (green), chloroplast autofluorescence (blue), and plant cell walls stained with propidium iodide (pink). Full and empty white triangles highlight epiphyllous and penetrated hyphae, respectively. Scale bar: 20 µm.

**Supplementary Figure S3. Recognition of AvrStb6 hinders the penetration of** Z. tritici**. A-F**. Confocal microscopy images of wheat cultivar Chinese Spring sprayed with *Z. tritici* 3D7-GFP and 3D7-GFP expressing AvtStb6_1E4_ (3D7-GFP+AvrStb6_1E4_). **A,B.** At 6 days post-infection (dpi, A), the virulent strain 3D7 penetrated the leaf and started invading its apoplastic space. At later stages (10 dpi, B), 3D7 invaded the leaf apoplast and accumulated in sub-stomatal cavities. **C-F**. 3D7-GFP reporter line ectopically expressing the *AvrStb6*_1E4_ gene does not penetrate through the stomata of Chinese Spring plants at 6 dpi (C, E) and at 10 dpi (D, F). Images of two independent transformant lines (Line A1 shown in C and D, and line A2 shown in E and F) depict identical observations. **G-L**. Confocal microscopy images of *Z. tritici* cytosolic reporter lines 1E4 (1E4-GFP) sprayed on Bobwhite and Bobwhite expressing the wheat resistance gene, *Stb6*. At 6 and 9 dpi, 1E4 penetrated and colonized the apoplast of Bobwhite (**G, H**). However, it remained at the leaf surface when inoculated on the Bobwhite expressing *Stb6* (**I-L**). Images of two independent wheat transformant lines (T2-19-2 shown in I and K, and T2-41-4 shown in J and L) depict identical observations at both 6 and 9 dpi. Rare instances of penetration could be observed, but those events present very limited apoplastic hyphal growth (J). Images A-L are maximum projections, and A’-L’ are orthogonal views. Dashed lines indicate the location of the orthogonal view on the corresponding image. Images are overlays of GFP signal (green), chloroplast autofluorescence (blue) and plant cell walls stained with propidium iodide (pink). Full and empty white triangles highlight epiphyllous and penetrated hyphae, respectively. Scale bar: 20 µm.

**Supplementary Figure S4. AvrStb6 accumulates in hyphal cells in direct contact with stomata. A-D**. Microscopy images of Bobwhite wheat plants infected with *Z. tritici* reporter line expressing AvrStb6_1E4_-GFP (3D7-mCherry+ AvrStb6_1E4_-GFP) at 6 and 9 days post-infection (dpi; A-B and C-D, respectively). In epiphytic hyphae, the GFP signal can only be observed in hyphal cells in direct contact with stomata (A and C). At both 6 and 9 dpi (B and D), hyphae that managed to penetrate through the stomata and colonize the leaf apoplast accumulate GFP. **E-J**. Bobwhite near-isogenic lines expressing *Stb6* (T2-19-2 and T2-41-4) were used to observe the AvrStb6_1E4_-GFP localization in the context of an incompatible interaction. At 6 and 9 dpi (E-G and H-J, respectively), the AvrStb6_1E4_-GFP signal was observed in hyphal in direct contact with stomata (E-J). Rare cases of hyphal penetration could be observed (F). In that case, hyphae presented a faint signal of AvrStb6 accumulation (F’). Images A-F are maximum projection overlays of GFP signal (green), mCherry signal (red) and chloroplast autofluorescence (blue). Images on A’-F’ are maximum projection overlays of only GFP signal (green) and chloroplast autofluorescence (blue). **K,L.** Microscopy images illustrating *AvrStb6*_1E4_ expression pattern. The 3D7 mTurquoise-labelled *Z. tritici* strain (in red), expressing a tagged histone protein (His1-mCherry) under the control of *AvrStb6*_1E4_ promoter, was sprayed on Runal wheat leaves. Fungal nuclei accumulating His1-mCherry signal (in green) indicate cells exhibiting *AvrStb6*_1E4_ promoter activity. Images were acquired at 6 and 9 dpi (K and L, respectively). Epiphytic hyphae show low expression levels of *AvrStb6*, except when the cells approach the stomata. In contrast, apoplastic hyphae strongly express *AvrStb6*. Full and empty white triangles highlight epiphyllous and penetrated hyphae, respectively. Scale bar: 20 µm.

**Supplementary Figure S5. Detection of AvrStb6** _1E4_ **protein produced in** Pichia pastoris. AvrStb6_1E4,_ coupled with Myc and His tags, was produced in *P. pastoris* and purified using Ni-NTA agarose beads. **A**. Western blot analysis of the AvrStb6_1E4_-Myc-His elutions fractions, separated by SDS-PAGE, and detected using an anti-His antibody. **B**. Coomassie Blue staining (left) and anti-His western blot detection (right) of the purified AvrStb6_1E4_-Myc-His, after Buffer exchange. Lane1: Precision plus protein ladder (5µl). Lane 2: AvrStb6_1E4_-Myc-His (20µl). The expected protein size is 18,5kDa.

**Supplementary Figure S6. Overall wheat gene expression pattern upon infiltration and spray infection. A.** Multidimensional scaling (MDS) plots of the expression data of 36 wheat RNASeq samples. The different colors correspond to different treatments, and the different shapes to different timepoints. **B.** MDS plot of the expression data of Chinese Spring leaves infiltrated with AvrStb6 protein and PBS buffer at three different timepoints (1 hours post infiltration (hpi), 2 hpi and 5 hpi). The different colors correspond to different treatments (dark blue = infiltrated with PBS buffer, dark orange = infiltrated with the AvrStb6 protein), and the three timepoints, 1, 2 and 5 hours post infiltration (hpi), are represented with different shapes. **C.** MDS plot of the expression data of Chinese Spring infected with the virulent strain 3D7, the avirulent strain 3D7+AvrStb6_1E4_ or mock at 3 or 6 days post infection (dpi). The different colors correspond to different treatments (light blue = mock, green = infected with 3D7 and orange = infected with 3D7+AvrStb6_1E4_) and the two timepoints are represented with different shapes.

**Supplementary Figure S7. Gene Ontology enrichment analysis for the identified sets of differentially expressed genes.** The “Biological processes” GO terms for which an enrichment has been observed in any of the gene sets are listed on the y-axis and the x-axis corresponds to the fold enrichment. The size and the color of the circles correspond to the number of genes and to the p-value respectively. GO terms were classified in the following categories: “plant hormones”, “photosynthesis, photorespiration, light stimulus and pigments”, “cell wall”, “stress responses”, “secondary metabolism”, “cell division, DNA and primary metabolism”, and “others”.

**Supplementary Figure S8. Gene Ontology enrichment analysis in “Molecular Functions” for the identified sets of differentially expressed genes.** “Molecular Functions” GO terms for which an enrichment has been observed are listed on the y-axis and the x-axis corresponds to the fold enrichment. The size and the color of the circles correspond to the number of genes and to the p-value, respectively.

**Supplementary Figure S9. Gene Ontology enrichment analysis in “Cellular Components” for the identified sets of differentially expressed genes.** “Cellular Components” GO terms for which an enrichment has been observed are listed on the y-axis and the x-axis corresponds to the fold enrichment. The size and the color of the circles correspond to the number of genes and to the p-value, respectively.

**Supplementary Table S1.** *Zymoseptoria tritici* strains used.

**Supplementary Table S2.** List of primers used for cloning and quantitative PCR.

**Supplementary Table S3.** RNASeq read counts statistics. For the 36 RNASeq samples, the percentage of reads mapping to the corresponding CDS as well as the number of reads mapping to the corresponding CDS are indicated.

**Supplementary Table S4.** Differential gene expression data of all the differential expressed genes in wheat. Gene Length is indicated in column B, counts are indicated for each replicate in columns C to AL. For each pairwise comparison, one column indicated the logFC and one the FDR adjusted p-value (from edgeR). For the infiltration experiment PBS vs AvrStb6 i) at 1hpi in columns AM and AN, ii) at 2hpi in columns AO and AP, iii) at 5hpi in columns AQ and AR. For the spray inoculation mock vs 3D7 iv) at 3dpi in columns AS and AT, v) at 6dpi in columns AU and AV; and mock vs 3D7_AvrStb61E4 vi) at 3dpi in columns AW and AX and vii) at 6dpi in columns AX and AZ. In columns BA to BG, the genes that are differentially expressed in each pairwise comparisons are indicated with "DE". Genes that are part of "DEGs in infiltration", "CORE DEGs", "3D7-specific DEGs" and "3D7_AvrStb61E4" are indicated in columns BH, BI, BJ and BL respectively. Previously cloned resistance genes Stb6, Stb15 and Stb16q are written in red, green and blue respectively.

**Supplementary Table S5.** Differential gene expression summary. The number of differential expressed genes identified with edgeR and DESeq2 are indicated. The overlap between the two methods has been considered for further analyses.

**Supplementary Table S6.** List of the biological processes GO terms enriched in enrichment analyses. The GO terms have been classified into categories indicated in columns 3 and 4.

**Supplementary Table S7.** Differential gene expression data of wheat homolog genes related to stomatal closure. Pairwise comparisons are indicated in columns C to P. For each pairwise comparison, one column indicated the logFC and one the FDR adjusted p-value. In columns Q to W, the genes that are differentially expressed in each pairwise comparisons are indicated with "DE". Genes that are part of "DEGs in infiltration", "CORE DEGs", "3D7-specific DEGs" and "3D7_AvrStb61E4" are indicated in columns X, Y, Z and AA, respectively.

**Supplementary Table S8.** List of the 247 genes responding specifically to the presence of AvrStb6. Superfamilies for which a hit was obtained when the protein sequence of the gene was blast against the conserved domain database of NCBI are listed in the column "Conserved Protein Domain Family".

**Supplementary Table S9.** List of the upregulated DEGs among the 247 “AvrStb6-specific genes”, their expression levels and conserved associated domains. List of upregulated genes upon infection with the avirulent strain at 3 dpi considering log2FC >1.5 and FDR <0.01 as shown in the colums J and K respectively (Comp_mock_Avr_3d). Espression levels for the other pairwise comparisons are also shown. infiltration experiment PBS vs AvrStb6 at 1hpi, 2hpi, and 5hpi. For the spray inoculation, mock vs 3D7 at 3dpi and 6dpi and mock vs 3D7_AvrStb61E4 at 3dpi and 6dpi are included. The following columns are the predicted functions and conserved domains. PANTHER_Family/Subfamily, Protein names UniProt, Protein families, conserved domains, hypothetical functions of each conserved domain, and protein function prediction after putting together the above information. NA= not assigned. Genes highlighted in light green correspond to the ones potentially involved in plant defense response (Supplementary Table S10). Genes highlighted in orange correspond to up-regulated genes encoding proteins involved in cell wall modification. Genes highlighted in blue potentially encode proteins playing a role in stomatal movement, sugar transporter protein (STP), and plant glutamate receptor (GLR).

**Supplementary Table S10.** Expression levels and protein domain predictions of upregulated genes among the 247 “AvrStb6-specific genes” encoding for proteins involved in plant defense response. For each pairwise comparison, one column indicated the logFC and one the FDR adjusted p-value (from edgeR). For the infiltration experiment PBS vs AvrStb6 i) at 1hpi, ii) at 2hpi, iii) and at 5hpi. For the spray inoculation mock vs 3D7 iv) at 3dpi, v) at 6dpi; and mock vs 3D7_AvrStb61E4 vi) at 3dpi and vii) at 6dpi. All genes in this table are differentially expressed. Genes that are part of "DEGs in infiltration", "CORE DEGs", "3D7-specific DEGs" and "3D7_AvrStb61E4" are indicated with a yes. Functional classification according to PANTHER, Uniprot and conserved domains (CDD, NCBI) are also indicated. The last column shows the biological function assigned to each protein. Abbreviations. RLK: receptor-like kinase; LRR: Leucine-rich repeat; RLP: Receptor-like protein; WAK: Wall-associated kinase; MLO: Mildew resistance locus O; NLR: Nod-like receptor. Genes highlighted in dark green correspond to the ones plotted in Figure 5C.

**Supplementary Table S11.** Differential gene expression data of all the differential expressed genes in *Z. tritici*. Gene Length is indicated in column B, counts are indicated for each replicate in columns C to N. For each pairwise comparison, one column indicated the logFC and one the FDR adjusted p-value (from edgeR). For the comparisons between isolates (3D7 versus 3D7AvrStb6_1E4) i) at 3dpi in columns O and P and ii) at 6dpi in columns Q and R. For the comparisons between timepoints (3dpi versus 6dpi) iii) in 3D7 in columns S ant T and iv) in 3D7_AvrStb61E4 in columns U and V. In columns W to Z, the genes that are differentially expressed in each pairwise comparisons are indicated with "DE". Genes that contain a signal peptide according to SignalP are listed in column AA and the ones that are effectors according to EffectorP in column AB.

